# Establishment and characterization of a new *Pseudomonas aeruginosa* infection model using 2D airway organoids and dual RNA sequencing

**DOI:** 10.1101/2023.03.11.532178

**Authors:** Cayetano Pleguezuelos-Manzano, Wouter A. G. Beenker, Gijs J.F. van Son, Harry Begthel, Gimano D. Amatngalim, Jeffrey M. Beekman, Hans Clevers, Jeroen den Hertog

## Abstract

*Pseudomonas aeruginosa* is a Gram-negative bacterium that is notorious for infections in the airway of cystic fibrosis (CF) subjects. Often, these infections become chronic, leading to higher morbidity and mortality rates. Bacterial quorum sensing (QS) coordinates the expression of virulence factors and the formation of biofilms at a population level. QS has become the focus of attention for development of alternatives to antimicrobials targeting *P. aeruginosa* infections. However, a better understanding of the bacteria-host interaction, and the role of QS in infection, is required. In this study, we set up a new *P. aeruginosa* infection model, using 2D airway organoids derived from healthy and CF individuals. Using dual RNA-sequencing, we dissected their interaction, focusing on the role of QS. As expected, *P. aeruginosa* induced epithelial inflammation. However, QS signaling did not affect the epithelial airway cells. The epithelium influenced several infection-related processes of *P. aeruginosa*, including metabolic changes, induction of type 3 and type 6 secretion systems (T3SS and T6SS), and increased expression of antibiotic resistance genes, including *mexXY* efflux pump and several porins. Interestingly, the epithelium influenced the regulation by QS of the type 2 (T2SS) and T6SS. Finally, we compared our model with *in vivo P. aeruginosa* transcriptomic datasets, from samples directly isolated from the airways of CF subjects. This shows that our model recapitulates important aspects of *in vivo* infection, like enhanced denitrification, betaine/choline metabolism, increased antibiotic resistance, as well as an overall decrease of motility-related genes. This relevant infection model is interesting for future investigations, helping to reduce the burden of *P. aeruginosa* infections in CF.

## Introduction

*Pseudomonas aeruginosa* is a Gram-negative opportunistic bacterium, which is known to chronically infect the airways of people with cystic fibrosis (CF). CF is a genetic disorder caused by mutations in the gene coding for the cystic fibrosis transmembrane conductance regulator (CFTR) protein. Mutated CFTR leads to an osmotic misbalance of the epithelial surface in multiple organs^1–3^. However, the airways are most affected due to their obstruction by desiccated mucus, which can lead to pulmonary failure^4,5^. This CF mucus presents a perfect condition for *P. aeruginosa* growth^6^. Because *P. aeruginosa* has a high intrinsic resistance to antibiotics, infections often become chronic^7–11^, contributing to the high morbidity and mortality observed in people with CF^10^.

A growing field of research focuses on quorum sensing (QS) as an alternative target to treat *P. aeruginosa* infections. Via QS, bacteria regulate a broad range of cellular processes based on their local cell density. QS in Gram-negative bacteria is highly conserved: a LuxI-type synthase produces a signal molecule (an acyl-homoserine lactone (AHL)) that can diffuse across membranes and bind to its cognate LuxR-type receptor, altering the expression of target genes^12,13^. Thus, when bacterial density is high, the concentration of AHLs increases, inducing downstream processes like biofilm formation and the production of virulence factors^12^.

*P. aeruginosa* presents a relatively complex QS network, involving three different systems: two typical LuxI/R systems (*las-*encoded and *rhl-encoded* system) and a unique Pseudomonas Quinolone Signal (PQS)-based system. These systems are hierarchical and highly interconnected. The Las system is the QS master regulator that induces the expression of the Rhl and PQS systems^14^. Importantly, inhibition of QS reduces the toxicity of *P. aeruginosa* in animal models and leads to faster clearing and prolonged survival of the infected animal^15–17^. However, the exact role of QS in human infection and CF is still underexplored^18^.

To date, *P. aeruginosa* CF infection models vary from the study of *P. aeruginosa* isolates from CF subjects^19–21^ to growing *P. aeruginosa in vitro* in artificial CF sputum and other bacterial media^22–26^, using various CF animal models^27–29^, working with *ex vivo* CF lungs^30,31^, or co-culture systems using cancer cell lines and primary cells ^32–37^. Each of these models present distinct advantages and disadvantages^38^.

Human airway organoids derived from adult stem cells faithfully resemble the cellular composition and physiology of the airway^39^. Additionally, airway organoids derived from CF subjects capture the molecular characteristics of the disorder and therefore are a useful tool to study CF phenotypes *in vitro^40^*. Furthermore, organoid co-cultures have recently been used to investigate the role of bacteria in colorectal cancer^41,42^. In the current study, we describe a new co-culture method using upper airway organoids grown in 2D-derived from healthy individuals and CF subjects-to study *P. aeruginosa* infections. By performing dual RNA-seq^43^, this analysis captures the interaction between the host cells and the bacteria.

## Materials and methods

### Bacterial strains and growth conditions

For this study, we used *P. aeruginosa* PAO1 strains, which constitutively express GFP. Stocks were stored in −80 °C in 20 % glycerol solutions. PAO1 strains were plated on Luria agar (LA) at 37 °C and grown in medium specific for the assay. *E. coli* RHO3 strains were used for conjugation and medium was supplemented with 400 μg/mL 2,6-Diaminopimelic acid (Sigma-Aldrich, Merck Life Science, Amsterdam, The Netherlands) to support growth. The PAO1 Δ*pqsA* strain has been described before^44^. The PAO1 Δ*lasI*/Δ*rhlI* strain was generated using allelic exchange following the method described before^45^. For the generation of the mutants we inserted upstream (using UP.Fw and UP.Rv primers) and downstream (using DN.Fw and DN.Rv primers) regions of the gene of interest in pEX18Gm plasmids using Gibson assembly restriction cloning, using the restriction enzymes SacI and Sph1 (Primers and plasmids are listed in Supplementary table 1 and 2). After Gibson assembly, the plasmid was transformed into RHO3 *E. coli* donor strains before conjugation via puddle mating with PAO1 GFP strain. Mutant colonies were identified with colony PCR (using seq.FW and seq.Rv primers) and confirmed by sequencing (performed by Macrogen Europe BV).

### Organoid cultures

Nasal brushing-derived epithelial stem cells were collected and stored with informed consent of all donor and was approved by a specific ethical board for the use of biobanked materials TcBIO (Toetsingscommissie Biobanks), an institutional Medical Research Ethics Committee of the University Medical Center Utrecht (protocol ID: 16/586). Nasal epithelial stem cells were isolated and expanded in 2D cell cultures as previously described^46^. After initial expansion, nasel cells were grown as organoids and cultured as previously described^39^. In brief, organoids were cultured in expansion medium containing Advanced DMEM F12, 1X GlutaMax (Life Technologies; 12634-034), 10mM HEPES (Life Technologies; 15630-056) (AdvDMEMF12++), supplemented with penicillin and streptomycin (10,000□IU/ml each; Life Technologies; 15140-122) 1× B27 supplement (Life Technologies; 17504-044), 1.2□mM N-acetyl-l-cysteine (Sigma-Aldrich; A9165), 10□mM nicotinamide (Sigma-Aldrich; N0636), 500□nM A83-01 (Tocris; 2939), 5□μM Y-27632 (Abmole; Y-27632), 1□μM SB202190 (Sigma-Aldrich; S7067), 100□ng/ml human FGF10 (PeproTech; 100-26), 25□ng/ml FGF7 (PeproTech; 100-19), 1% (vol/vol) RSPO3, and Noggin (produced via the r-PEX protein expression platform at U-Protein Express BV). For its passage, organoids were collected, washed and resuspended in TrypLE (Gibco) and incubated at 37°C for 15 minutes. Then, organoids were mechanically disrupted into single cells and plated in droplets of Cultrex growth factor reduced BME type 2 (Biotechne | R&D systems 3533-010-02). For the 2D culture of airway organoids, 10^5^ cells were seeded into 24-well polystyrene membranes (Greiner Bio-One) and cultured for one week in expansion medium in both top and bottom compartments until confluency. After confluency, cells were differentiated in air liquid interface during 1 month using PneumaCult™-ALI Medium (Stem cell technologies) supplemented with Hydrocortisone stock solution (5 μl/ml; Stem cell technologies #07925) and Heparin solution (2 μl/ml; Stem cell technologies #07980).

### 2D airway organoid - *Pseudomonas aeruginosa* co-culture

Bacterial colonies were picked and grown in DMEM medium (ThermoFisher Scientific, 31966-021), supplemented with 10mM HEPES (ThermoFisher Scientific, 15630080) and 1X GlutaMAX (ThermoFisher Scientific, 35050061) until an OD_600_ = 0.35 – 0.45. 1 mL of early log-phase bacterial culture was taken and centrifuged for 3 min at 15,000 g, before washing once with PBS. Bacterial cells were pelleted again and resuspended in AdvDMEMF12++ to normalize to OD_600_ = 0.4. Bacteria were diluted 100 x in the same medium before adding 50 μL to the organoids to reach a multiplicity of infection (MOI) of 0.1. Organoids in transwells were washed three times in AdvDMEMF12++ (without antibiotics) before addition of the bacteria. Co-cultures were incubated at 37°C with 5 % CO_2_ for 14 h. In parallel, 50 μL of bacteria were grown in a 96-well plate and organoid transwells without bacteria were incubated as controls. The epithelial cells were checked for damage using the fluorescent EVOS FL Auto 2 microscope (ThermoFisher Scientific) at 14 h. Damaged co-cultures were excluded for analysis.

### Imaging of organoid 2D cultures and PAO1-GFP co-culture

Organoid 2D culture and PAO1 co-cultures were fixed with 4% formaldehyde for 2 hours at room temperature. Then, 2D cultures were processed for immunohistochemistry following standard techniques and embedded in paraffin. After sectioning, hematoxicilin/eosin staining was performed according to manufacturer instructions. After blocking, the used primary antibodies included anti-MUC5AC (Thermo, MA5-12175), anti-acetylated Tubulin (SantaCruz, sc-23950) and anti-P63 (Abcam, ab735). Co-cultures were processed for wholemount immunofluorescence 41 imaging using standard techniques^41^. Then, co-culture was stained with Phalloidin Atto-647 (65906-10NMOL) and DAPI and imaged with SP8 confocal microscope (Leica).

### Colony forming unit (CFU) test

For the CFU test, organoid infection protocol was followed as described before. For time point 0 h, plates were incubated for only 5 min. For time point 14 h, plates were incubated 14 h. After incubation, 50 μL of 0.5% saponin was added to the wells and incubated for 10 min at room temperature. Then, the volume was resuspended and transferred to Eppendorf tubes before centrifugation for 10 min at 15,000 g. Supernatant was aspirated and pellets were resuspended in PBS before making 10-fold dilutions. 5 μL of the dilutions were plated on tryptic soy agar and plates were incubated overnight at 34 °C before the colonies were counted.

### RNA isolation

RNA was isolated from the cultures using the MasterPure Complete DNA and RNA Purification Kit (Immunosource, Belgium) following manufacturer instructions. To remove all the gDNA, two additional rounds of DNase treatment were performed using TURBO DNase (ThermoFisher Scientific) following manufacturer’s protocol. RNA was subsequently isolated again using the MasterPure RNA isolation kit and the RNA was dissolved in MQ and stored at −80 °C until further use. For RNA-sequencing, RNA libraries were validated with Agilent 2100 bioanalyzer. Samples were sent for sequencing to the Utrecht Sequencing Facility (Useq), for library preparation using Truseq RNA stranded ribo-zero. RNAseq was performed using the Illumina NextSeq2000 platform.

### Mapping of raw-reads to the genome

To analyze the dataset, a variety of bioinformatics tools were used and we followed a similar protocol as described before, with minor modifications (Supplementary figure 1)^47^. First, the reads were mapped to the human genome (hg19) using STAR mapping 48 software^48^. Unmapped reads were written into a new FastQ-file. Gene counts were assigned to the mapped reads using FeatureCounts^49^. Unmapped reads were then mapped against the PAO1 genome (NCBI: txid208964)^48,50,51^. Gene counts were again assigned using FeatureCounts. The code for this pipeline is available on Github (https://github.com/GJFvanSon/Hubrecht_clevers.git).

### Bioinformatic analysis

Human and PAO1 count tables were independently analyzed using DESeq2^52^ DEGs were calculated using the lfcShrink function with argument type set to “apglm”. Volcano plots were generated using EnhancedVolcano package^53^. For the human dataset, GO enrichment analysis was performed using the function enrichGO() from the package clusterProfiler()^54^ with arguments OrgDb=org.Hs.eg.db, ont=“BP”, pAdjMethod=“fdr”, minGSSize=“1”, maxGSSize=“2000”, qvalueCutoff= “0.05” and readable=“TRUE”. For PAO1 dataset, GO enrichment analysis was performed using the online tool PANTHER 17.0, with the following settings: analysis type: PANTHER overrepresentation test (Released 20221013), annotation version and release date: GO ontology database DOI: 10.5281/zenodo.6799722 released 2022-07-01, reference list: *Pseudomonas aeruginosa* (all genes in database), annotation data set: GO biological process complete, test type: Fisher’s exact, correction: calculate false discovery rate. KEGG pathway was generated using the Pathview R package. Pseudomonas category and GO gene lists were downloaded from *Pseudomonas* Genome DB version 21.1 (2022-11-20)^55^. PCA were plotted using the DESeq2 function plotPCA(). ntop parameter was set to 1500 for PCA of *in vivo* samples integration. Protein-protein association network analysis was performed using the STRING v11 online tool^56^. Common genes with a log2Foldchange / logSE > |4| were used as input. The input parameters were set as follow: Organism = “Pseudomonas aeruginosa PAO1”, Network Type = “full string network”, Required score = “high confidence (0.700)”, and FDR stringency = “medium (5 percent)”. MCL clustering was performed on the resulting network using the following parameters: inflation parameter = “3”, and edges between clusters = “Don’t show”.

## Results

### Establishment of 2D airway organoid co-cultures with *P. aeruginosa*

We aimed to study the interaction between *P. aeruginosa* and the airway epithelium during early infection. For this, we established an epithelial co-culture system composed of upper airway (nasal) organoids cultured in 2D and the well-characterized *P. aeruginosa* PAO1 strain constitutively expressing GFP. Organoid cultures were differentiated for 1 month in 2D air liquid interface (ALI) (Figure 1A). This approach gave rise to a pseudostratified epithelium containing the main cell types of the mature airway epithelium: goblet cells marked by MUC5AC, ciliated cells marked by acetylated tubulin, and basal cells marked by TP63 (Figure 1B). 2D ALI cultures allow for easy apical exposure to bacteria. PAO1 bacteria were pipetted apically at a multiplicity of infection (MOI) of 0.1 (Figure 1C). After 14 hours, bacterial aggregates had formed on the epithelial cells implying that downstream pathways including biofilm formation are active in co-culture (Figure 1D, E). Longer co-culture time was not feasible due to elevated epithelial cell death caused by the bacteria. Wild type PAO1 cells and PAO1 strains ΔQS (*rhlI*/*lasI* KO) and Δ*pqsA* (*pqsA* knockout) that display impaired QS were alive and proliferated during this time span (Figure 1F).

**Figure 1.**
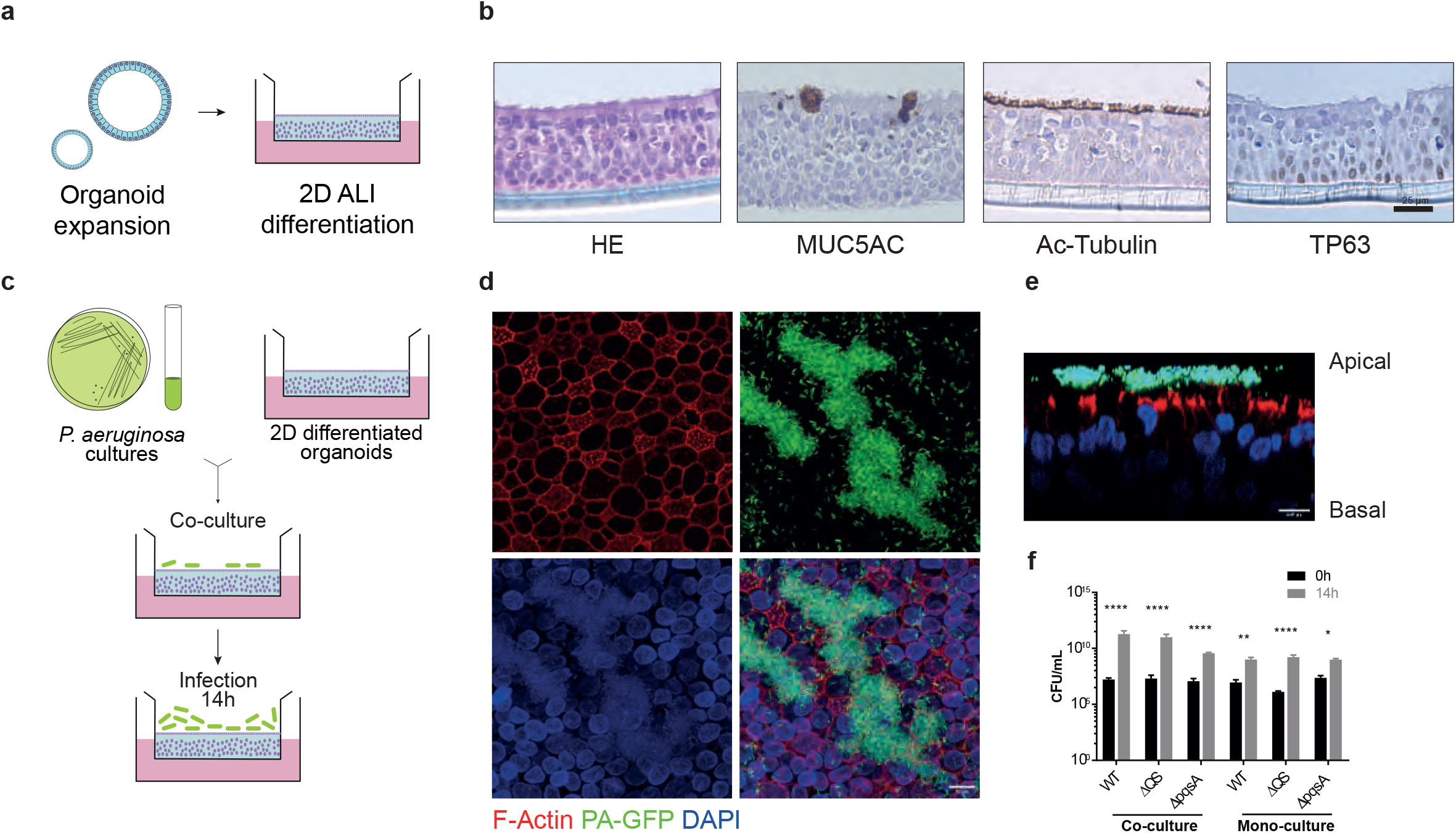
Co-culture establishment of 2D-airway organoids with *P. aeruginosa* PAO1. **a)** Schematic representation of organoid 2D culture establishment and ALI differentiation. **b)** HE and staining of the major cell types present in ALI differentiated airway organoids, goblet cells stained by MUC5AC, ciliated cells by acetylated (Ac) tubulin and basal cells by TP63. Scale bar indicates 25 μm **c)** Schematic representation of co-culture establishment of PAO1-GFP and differentiated airway organoids. **d)** Z projections and **e)** cross-section of confocal imaging of the co-culture after 14h. Red: FActin; green: PAO1-GFP; blue: DAPI. Scale bar indicates 10 μm. **f)** CFU assay of WT PAO1 bacteria and PAO1 ΔQS and Δ*pqsA* strains following co-culture with organoids and in liquid medium at time points 0h and 14 h. The mean of the triplicates was plotted and error bars represent standard error of the mean (SEM). To determine statistical significance between the time points, log-transformed data was analyzed using two-way ANOVA, corrected for multiple comparisons using Sidak’s test (*, P<0.05; **, P<0.005; ****, P<0.0001).

### Dual RNA-sequencing of the co-culture model

To study the interplay between upper airway epithelial organoids and *P. aeruginosa*, we subjected the 14 hours co-cultures to dual RNA-seq, in order to capture the transcriptomic response of both components. For this, we used organoid lines derived from a healthy donor or from a CF subject and either WT, Δ*pqsA* or ΔQS PAO1 cells (Figure 2A). Organoid cultures without bacteria and pure bacterial cultures were used as controls. As an initial step to validate the feasibility of our approach, we performed bacterial RNA sequencing of PAO1-WT strain, both in co-culture and by itself (Supplementary figure 1). We added these samples to the analysis of the dual RNA-seq cohort unless otherwise stated. After a two-step mapping approach (Supplementary Figure 1A), approximately half of the reads from co-culture samples were aligned to the bacterial or human genome, indicating that the dual RNA-seq approach efficiently captured the transcriptome of both components. As expected, the analysis of separately cultured 2D organoids and bacteria yielded almost 100% reads of the corresponding species (Figure 2B).

**Figure 2.**
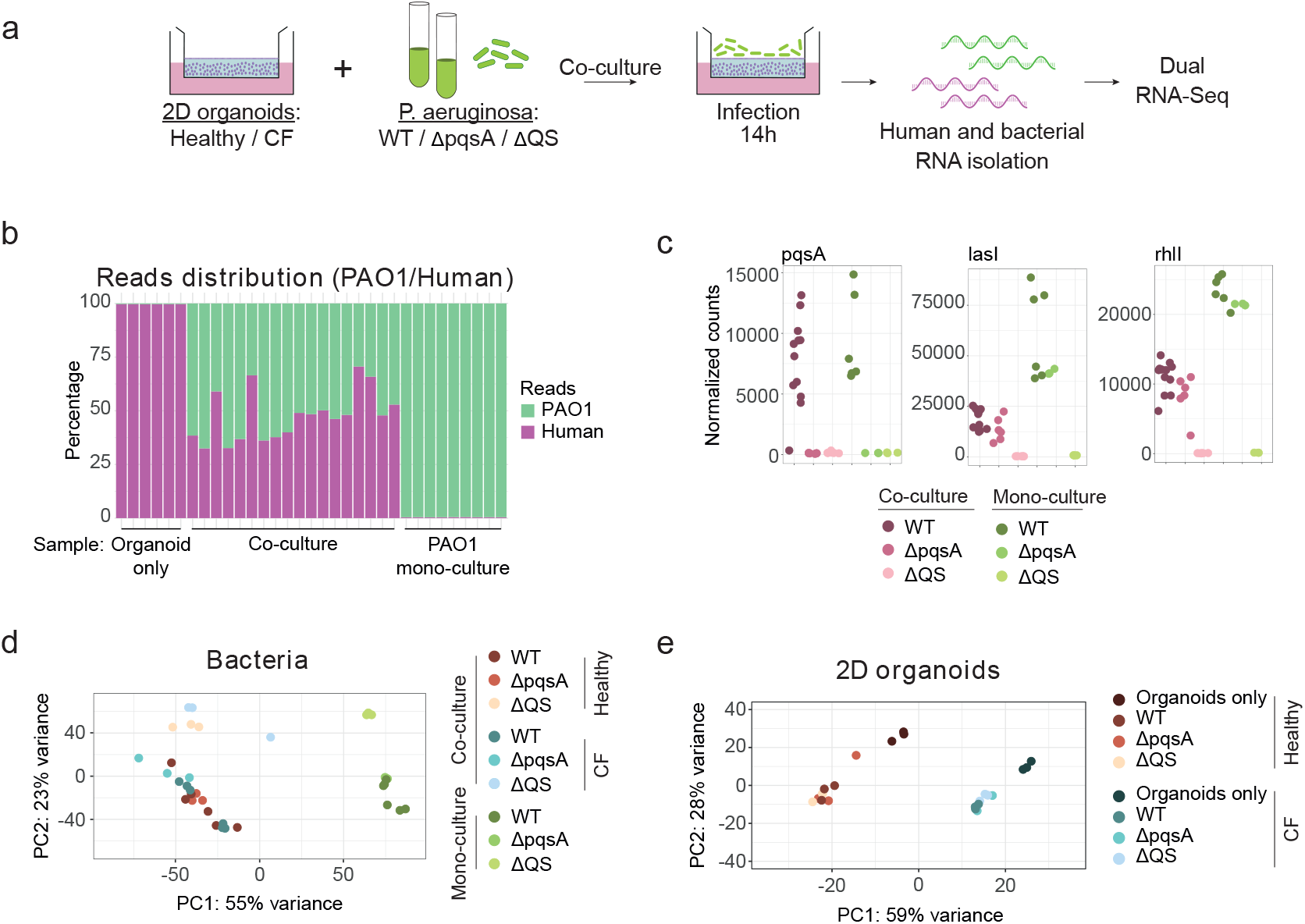
Co-culture characterization by Dual RNA-seq. **a)** Schematic representation of the Dual RNA-seq experiment. **b)** Distribution of human and bacterial reads across the different samples included in the run after performing the mapping and count assignment (as in Supplementary figure 1). **c)** Knock-out validation by gene expression. Normalized counts of *pqsA, lasI* and *rhlI* across the different samples of the cohort. Color code indicates culture condition (Green: mono-culture; magenta: co-culture) and PAO1 genotype (Dark: WT; middle: *ΔpqsA;* light: ΔQS). **d)** PCA plot of PAO1 samples. **e)** PCA plot of 2D organoid samples. Color code indicates PAO1 genotype, culture condition (co-culture or mono-culture) and organoid genotype (Healthy or CF).

We first validated the differential expression of the bacterial KO genes in the bacterial strains. As expected, the PAO1 Δ*pqsA* strain showed inhibition of *pqsA*, while not affecting *lasI* and *rhlI*. PAO1 ΔQS showed inhibition of all three QS pathways, including inhibition of *pqsA* due to the hierarchical QS system in *P. aeruginosa* PAO1 (Figure 2C). These three strains allowed us to study and compare the effect of QS and of the PQS system in co-culture. Comparison of the PAO1 transcriptomes showed a major difference between PAO1 cells grown in mono-culture and in co-culture with organoids (Figure 2D). The effect of ΔQS was also evident in co-culture and in bacterial monoculture (Figure 2D). However, Δ*pqsA* did not show a marked transcriptional change, other than *pqsA* expression itself, compared to WT bacteria, irrespective of the culture method (Figure 2D). Since our analysis included only one healthy and CF organoid line, conclusions about CF-specific effects of PAO1 cannot be drawn. Nevertheless, both organoid lines showed a clear transcriptional response to the presence of PAO1 irrespectively of the bacterial genotype (Figure 2E).

### *P. aeruginosa* induces epithelial inflammation

Next, we focused on the effect induced by the different PAO1 strains on the 2D organoid cultures (Figure 3A). Coculture with PAO1 cells led to upregulation of 1610 genes in 2D organoids and downregulation of 638 genes. This occurred irrespectively of the PAO1 genotype (Fig. 3B) For both healthy and CF 2D organoids, these changes reflected pathways involved in NF-κ□-mediated inflammation and response to lipopolysaccharides (LPS), including *IL1A/B*, *TNFA*, and various *CXCL* chemokines (Figure 3D and E). Whereas previous studies have shown that QS molecules affect epithelial cells^57–59^, no clear QS-derived effect was observed when organoids were exposed to PAO1 WT compared to Δ*pqsA* and ΔQS strains (Figure 3B). Since the Δ*pqsA* and ΔQS strains lack expression of some or most QS signaling molecules, this observation contrasted with a previous report suggesting that the epithelium can sense and respond to QS molecules^57^.

**Figure 3.**
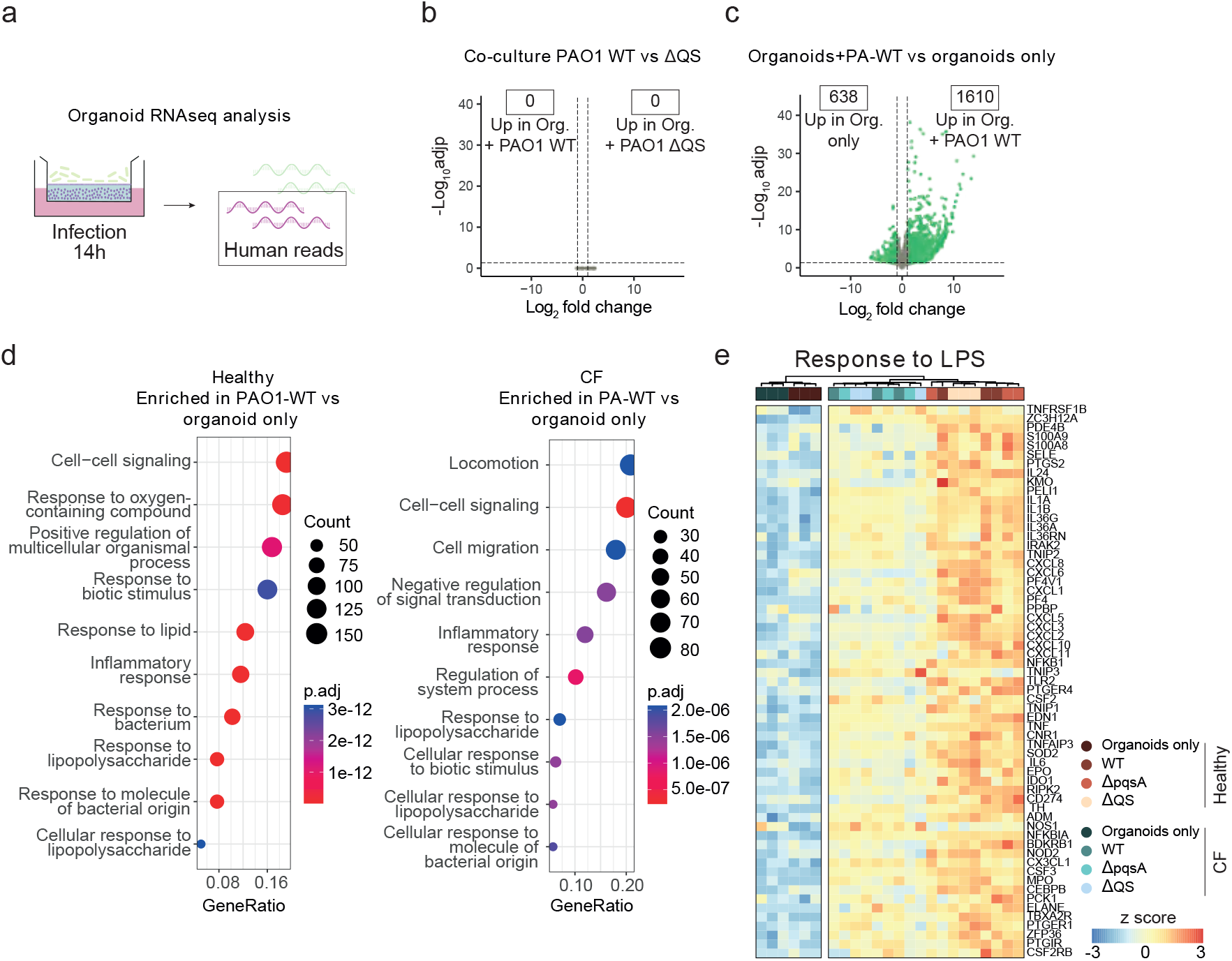
Transcriptional response of the epithelium to infection with the different PAO1 strains. **a)** Schematic representation of the analysis. **b)** Volcano plot showing the log_2_ fold change and −log_10_ adjusted p-value of all genes, when comparing the transcriptome of 2D organoids exposed to PAO1 WT or PAO1 ΔQS. **c)** Volcano plot showing the Log_2_ fold change and −log_10_ adjusted p-value per gene comparing the transcriptome of 2D organoids exposed to PAO1 WT or unexposed controls. Green indicates differentially expressed genes (DEGs) (log_2_ fold change > 1 and adjusted p value < 0.05). **d)** Gene ontology enrichment analysis showing top 10 categories enriched in 2D organoids exposed to PAO1 WT. Left panel: Healthy organoid line. Right: CF organoid line. **e)** Gene expression heat map of genes from “Response to lipopolysaccharide” GO term category (GO:0032496). Color code indicates culture condition (co-culture or mono-culture) and PAO1 genotype (WT, Δ*pqsA* or ΔQS).

### The epithelium induces *P. aeruginosa* transcriptional changes associated with infection

We next focused on the effect of the organoids on *P. aeruginosa* PAO1 (Figure 4A). Comparing the gene expression profile of PAO1 grown in co-culture versus monoculture revealed a total of 2215 differentially expressed genes (DEGs) (979 upregulated in co-culture; 1136 upregulated in mono-culture) (Figure 4B). Gene ontology enrichment analysis of these genes revealed broad metabolic differences between the two culture modes. Genes involved in iron acquisition (e.g., siderophore and pyoverdine processes) showed higher expression in pure bacterial cultures (Supplementary Figure 2A). This contrasted with what has been observed in human infections^19,60^, but was in agreement with previous co-culture attempts^35^.

**Figure 4.**
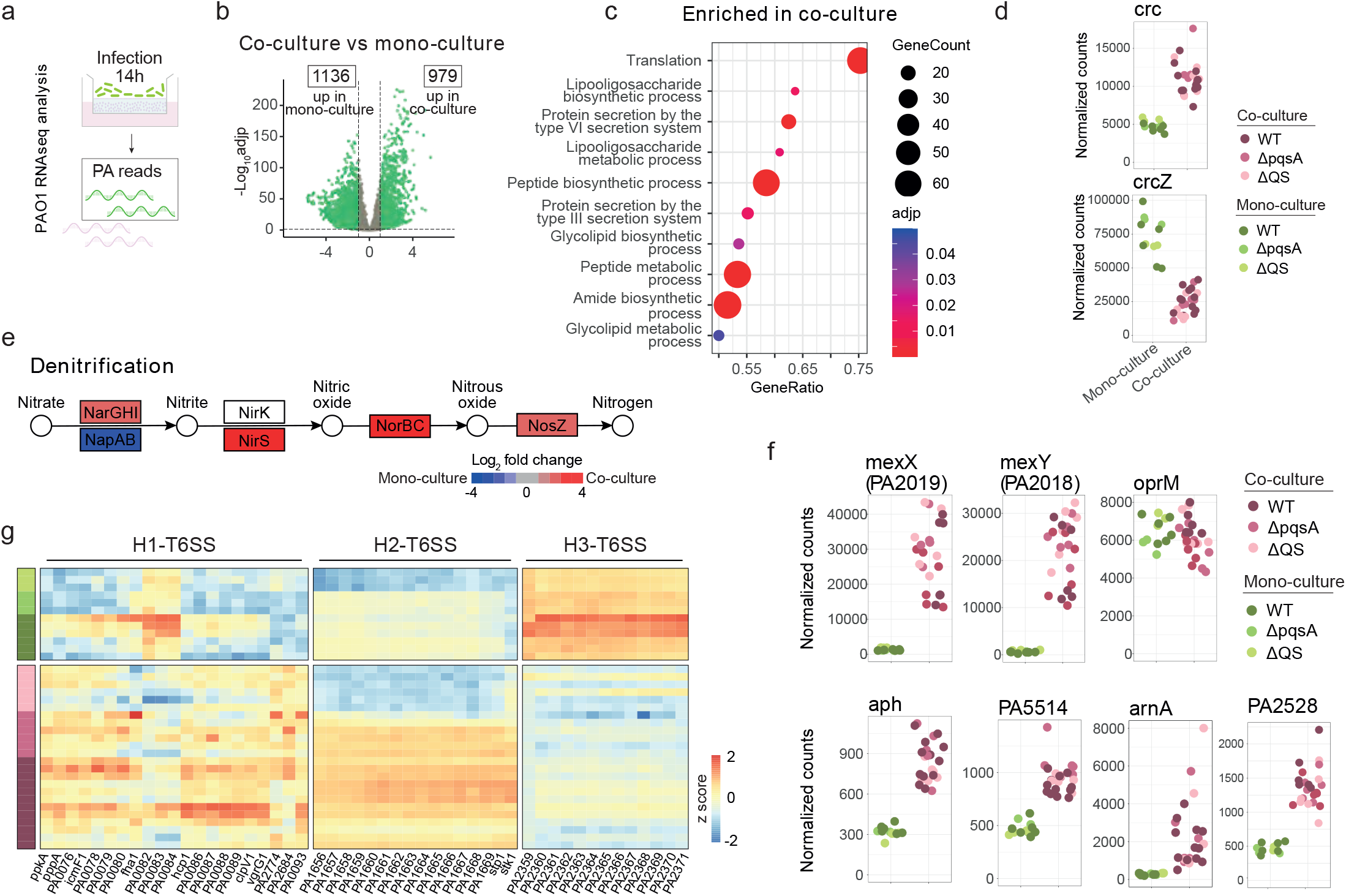
Transcriptional response of PAO1 to the presence of airway epithelium. **a)** Schematic representation of the analysis. **b)** Volcano plot displaying the Log_2_ fold change and −log_10_ adjusted p-value of all genes, when comparing the PAO1 transcriptomes of co-culture and bacterial mono-culture samples. Green indicates differentially expressed genes (DEGs) (log_2_ fold change > 1 and adjusted p value < 0.05). The number of genes upregulated in co-culture and bacterial mono-culture is indicated. **c)** Gene ontology enrichment analysis showing top 10 categories enriched in PAO1 exposed to airway epithelium in co-culture. **d)** Normalized count plots of genes involved in CCR pathway, *crc* and *crcZ*. **e)** KEGG pathway pae00910 plot displaying the log_2_ fold change of genes involved in denitrification. DEGs from co-culture vs bacterial culture mono-culture comparison of PAO1 transcriptomes. **f)** Normalized count plots of genes involved in *P. aeruginosa* antibiotic resistance. **g)** Heat map displaying expression of genes involved in *P. aeruginosa* T6SS. Genes grouped by H1, H2 or H3 T6SS subtype^68,69^. Samples grouped by culture condition. Color code indicates culture condition (Green: bacterial mono-culture; magenta: co-culture) and PAO1 genotype (Dark: WT; middle: Δ*pqsA;* light: ΔQS).

PAO1 cells co-cultured with airway cells increased their expression of genes related to peptide, glycolipid and amide biosynthetic pathways (Figure 4C), suggesting major metabolic rearrangements. Interestingly, co-cultured PAO1 cells presented increased *crc* and decreased *crcZ* levels (Figure 4D), two main regulators of the carbon catabolite repression (CCR) pathway^61,62^. This pathway is central to the hierarchical utilization of preferred carbon sources by *P. aeruginosa*. This finding suggested that the epithelium provides a source of preferred nutrients compared to the medium alone. Interestingly, expression change was observed in only a subset of genes known to be regulated by the CCR pathway (Supplementary Figure 2B). This highlights the complexity of metabolic regulation.

Another important aspect of *P. aeruginosa* infection of individuals with CF is its ability to perform denitrification^63^. This mechanism enables the utilization of nitrogenous oxides (nitrate, nitrite, and nitrous oxide) as electron acceptor for respiratory growth in anoxic conditions, such as during the course of infection^64^. We found that co-cultured PAO1 expressed increased levels of many genes involved in nitrogen metabolism (Supplementary Figure 2C), and particularly those used in denitrification (Figure 4E). This suggests that there is local anoxia due to the oxygen consumption by the epithelium and high-density bacterial population, which is similar as observed in airway infections^6,65^. Beyond metabolism, the bacteria also showed elevated levels of genes involved in resistance to antibiotics (Supplementary Figure 2D). Particularly striking was the effect of the epithelium on genes encoding the MexXY efflux pump, porins, and genes like *aph, PA5514, arnA* and *PA2528* encoding antibiotic degrading enzymes (Figure 4F and Supplementary Figure 2E). Of note, our co-culture system is performed in antibiotic-free conditions. Finally, the presence of epithelial cells induced the expression of type 3 (Supplementary Figure 2F) and type 6 (Figure 4G) secretion systems (T3SS and T6SS). These bacterial secretion systems are syringe-like structures used to inject toxins into the cytoplasm of target cells^66–69^. The epithelium mainly induced the expression of the H2-T6SS, and to a lesser extent H1-T6SS, but it repressed those belonging to the H3-T6SS subtype. H1 and H2 subtypes are known to act against other prokaryotes and eukaryotes, respectively^68,70^. Little is known about the role and regulation of H3-T6SS in infection.

### The epithelium influences aspects of *P. aeruginosa* QS regulation

Next, we investigated how the presence of the epithelium affected QS-regulated processes in *P. aeruginosa* PAO1. Since only the dual RNA-seq run contained samples from all three bacterial conditions (WT, *ΔpqsA*, and ΔQS), only samples from this run were included in the analysis to avoid batch-induced bias. In general, LasR- and RhlR-regulated genes and only some PQS-regulated genes were downregulated in PAO1 WT co-culture conditions compared to bacterial mono-cultures (Figure 5A). This correlates with previous descriptions of stronger QS-induced responses in pure bacterial cultures than in clinical infections^19,65,71^. Interestingly, the QS receptor levels (*mvfR, lasR* and *rhlR*) seemed to be more affected by ΔQS when the bacteria were co-cultured with the epithelium than in mono-culture. This could impact the QS regulatory network and therefore it is worth taking into consideration when interpreting results of QS regulation using *in vitro* models. Only 46 and 23 DEGs were found when comparing WT PAO1 with Δ*pqsA* in co-culture and in mono-culture respectively (Supplementary Figure 3A-B). From these, the *antABC* operon was affected only in mono-culture (Supplementary Figure 3C). This operon encodes the enzymes responsible for derivatizing the PqsA substrate anthranillic acid to catechol, before degradation to intermediates of the tricarboxylic acid (TCA) cycle^72^. The loss of PqsA in the Δ*pqsA* strain could lead to an accumulation of anthranillic acid. Via upregulation of *antABC* in bacterial mono-cultures, anthranillic acid might be used as a nutrient.

**Figure 5.**
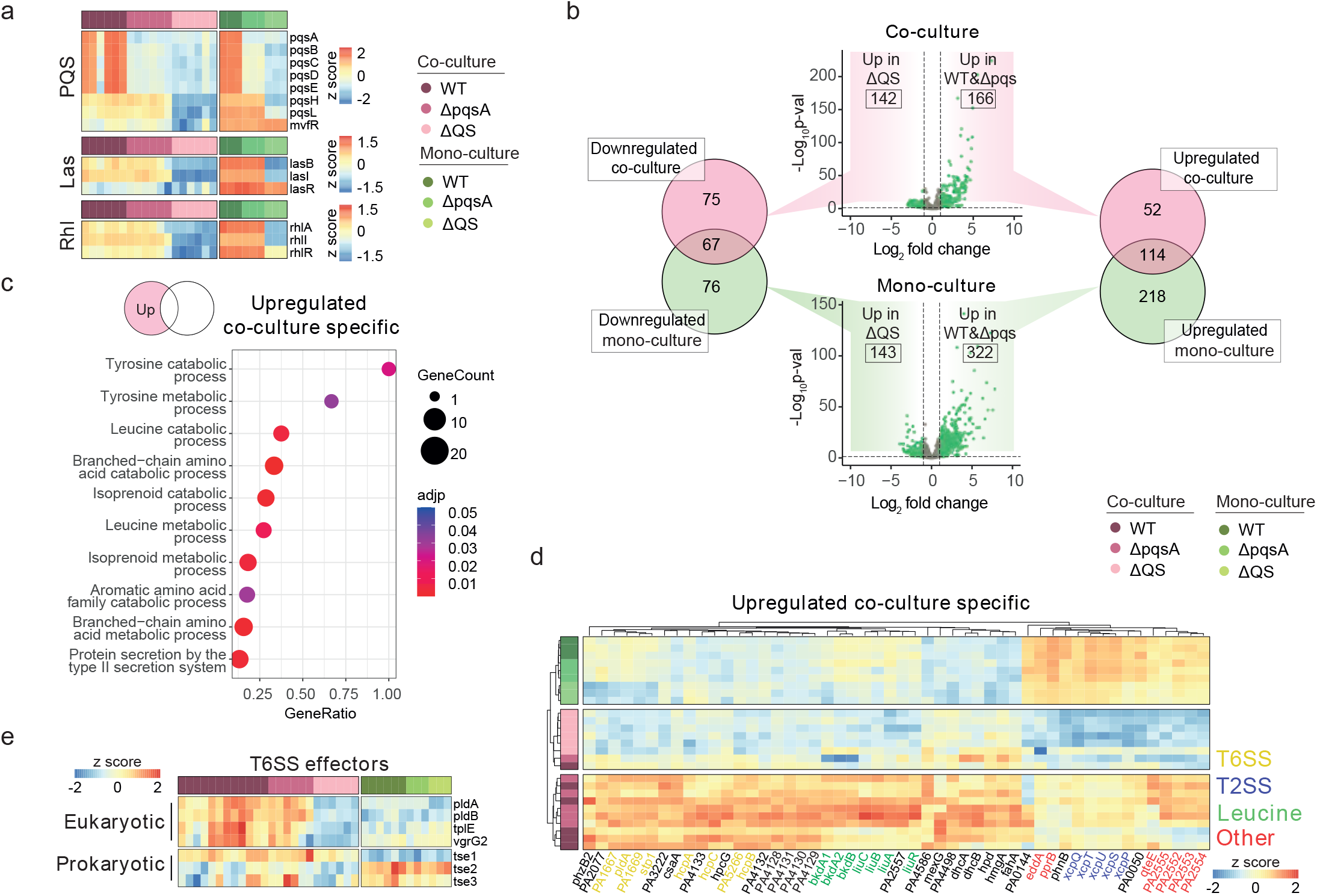
Epithelial effect on PAO1 QS regulation. **a)** Gene expression heat map of genes involved in PQS, Las or Rhl QS pathways. **b)** Volcano plots displaying gene log_2_ fold change and −log_10_ adjusted p-value when comparing the transcriptomes of WT and Δ*pqsA* PAO1 to those of ΔQS in co-culture (top) and in pure bacterial cultures (bottom). Venn diagrams display the overlap between genes up (right) and downregulated (left) in the comparisons. **c)** Gene ontology enrichment analysis showing top 10 categories enriched in genes that are specifically upregulated in co-culture in WT and Δ*pqsA* PAO1 transcriptomes compared to ΔQS. **d)** Gene expression heat map showing top 50 co-culture-specific DEGs. Genes are color-coded according to the following categories (Yellow: T6SS; purple: T2SS; green: Leucine metabolism; red: other pathways). **e)** Gene expression heat map of T6SS eukaryotic and prokaryotic effectors. Sample color code indicates culture condition (Green: bacterial culture mono-culture; magenta: coculture) and PAO1 genotype (Dark: WT; middle: Δ*pqsA;* light: ΔQS).

In contrast, deletion of *rhlI* and *lasI* in ΔQS led to 308 and 465 DEGs in co-culture and in mono-culture, respectively (Figure 5B). In order to dissect which QS-regulated processes were affected by the epithelium, we identified DEGs that occurred specifically in co-culture (52 up and 75 downregulated). Iron uptake pathways were downregulated only in ΔQS co-culture condition (Supplementary Figure 3D). This highlighted the differences of the two culture systems regarding iron utilization by the bacteria. On the other hand, genes involved in leucine and tyrosine catabolism were upregulated (Figure 5C-D). The amino acid utilization by *P. aeruginosa* is thought to be a key element of *P. aeruginosa* adaptation to the human airways, since amino acid auxotrophy is common in CF clinical isolates. This could explain why the QS effect is not observed in isolated bacterial cultures ^73^ Furthermore, we found genes regulated by QS specifically in coculture involved in bacterial adhesion and biofilm formation (*pprB*), and phosphatase and phosphodiesterase activities (*eddA*)^74,75^. Additionally, a number of T2SS and T6SS related genes (Figure 5C-D) were specifically regulated by QS in co-culture. T2SS-related genes from WT PAO1 in co-culture showed slightly reduced levels compared to mono-culture. In contrast, QS mutants in co-culture showed a greatly reduced expression of T2SS-related genes. This suggested that the epithelium potentially inhibits T2SS, which can be counteracted by QS-regulated molecules in WT bacteria. The expression of T6SS-related genes was low in all PAO1 strains in mono-culture. Coculture conditions specifically showed a QS-regulatory effect on the expression of these genes. This suggested that a combination of epithelial and QS factors induce the expression of some T6SS genes. In addition, T6SS effectors that are involved in pathogenicity of eukaryotic cells were both induced by a combination of epithelial and QS signals (Figure 5E).

### Benchmarking 2D co-culture model with chronic clinical samples

Next, we addressed which aspects from *in vivo P. aeruginosa* infections were recapitulated in our 2D co-culture model. We compared the transcriptional profiles of clinical *P. aeruginosa* strains directly isolated from airway biopsies from CF subjects to the PAO1 co-culture samples. Pure bacterial culture samples from this and other studies were included in the comparison^19,65,76^ (Supplementary Figure 4). After dataset integration, PCA analysis revealed that the origin of the bacterial samples (*in vivo*, coculture, or mono-culture) explained the clustering of the samples best (Figure 6A). The comparison of the *in vivo* and co-culture transcriptomes to all mono-culture samples revealed a total of 1382 and 2045 DEGs, respectively (Figure 6B). Despite the fact that our co-culture represented an early stage of infection compared to the chronic state of the clinical samples, 269 genes were common between both comparisons (4.7% of all PAO1 genes) (Figure 6C-D). This core gene signature captured the aspects of an *in vivo* infection present in our co-culture model. In order to understand pathways enriched in this core gene signature, we performed protein-protein interaction network analysis (Figure 6E). This confirmed that denitrification is an important process for *P. aeruginosa* infection and that this was captured by our model (Figure 6E-F, Figure 4E and Supplementary Figure 5). Increased expression of *mexYX* antibiotic efflux pump (Figure 6E-F, Figure 4F, Supplementary Figure 5) was also confirmed *in vivo*. Additionally, the expression of some T2SS proteins (Figure 6E-F, Supplementary Figure 5) was reduced in the core signature compared to *in vitro* cultures, which is in line with our previous analysis (Figure 5D).

**Figure 6.**
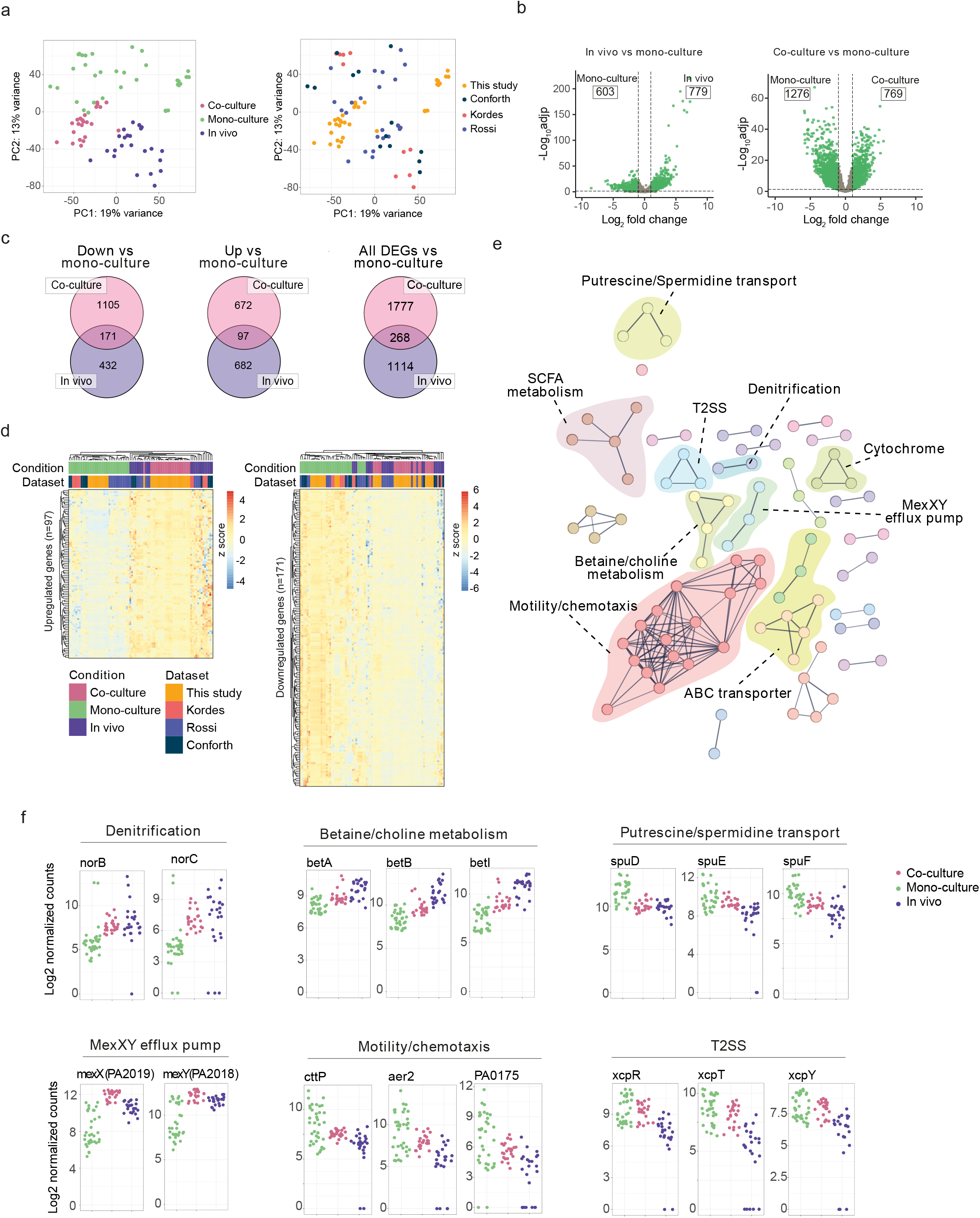
Benchmarking co-culture model with *in vivo P. aeruginosa* transcriptomic datasets directly isolated from the airways of CF subjects. **a)** PCA plots showing sample distribution by condition (Magenta: co-culture; green: bacterial culture in isolates; purple: *in vivo*) or by study of origin (Orange: this study; purple: Cornforth et al., 2018^19^; pink: Kordes et al., 2019^76^; blue: Rossi et al., 2018^65^). **b)** Volcano plots displaying gene log_2_ fold change and −log_10_ adjusted p-value comparing transcriptomes of *in vivo P. aeruginosa* (left) or co-cultured PAO1 (right) to those of all pure bacterial culture samples. Green indicates differentially expressed genes (DEGs) (log_2_ fold change > 1 and adjusted p value < 0.05). Indicated in the boxes the number of up-or downregulated DEGs. **c)** Venn diagrams displaying the overlap between genes that are upregulated (left), downregulated (middle), or both (right) in the previous *in vivo* and co-culture comparison to *in vitro* and mono-culture samples (b). **d)** Expression heat map displaying the common up-(left) and downregulated (right) genes. Samples clustered based on the expression of all genes plotted per heat map. Color-code indicates condition (Magenta: co-culture; green: pure bacteria; purple: *in vivo*) and study of origin (Orange: this study; purple: Cornforth et al., 2018; pink: Kordes et al., 2019; blue: Rossi et al., 2018). **e)** Protein-protein interaction network of common DEGs (in vivo and co-culture. Each node represents a protein encoded by a DEG. Edges represent known protein-protein association (either physical or functional) with a confidence level higher than 0.7. Node color represent clusters generated MCL method. Highlighted the pathway to which the cluster proteins belong. **f)** Log_2_ normalized count plots of representative genes from pathways highlighted by the network analysis. Colorcode indicates Magenta: co-culture, green: pure bacteria and purple: *in vivo*.

Beyond confirming the relevance of some of the pathways previously discovered by our cohort, the comparative analysis (Figure 6) uncovered other processes relevant for *in vivo* infection. This included elevated levels of choline/betaine metabolic genes (*betABI*), responsible for the production of glycine-betaine (GB) (Figure 6E-F, Supplementary Figure 5). The accumulation of GB has been proposed as a bacterial osmo-protective mechanism^65^, and to be important in *P. aeruginosa* infection in mice^77^. Additionally, our 2D co-culture model captured the reduced levels of motility- and chemotaxis-related genes that are observed *in vivo* compared to pure bacterial cultures (Figure 6E-F, Supplementary Figure 5), which is another important aspect for biofilm formation.

## Discussion

In this study, we describe a novel *P. aeruginosa* co-culture system using 2D human airway organoids derived from healthy and CF individuals. Subjecting the co-culture to dual RNA-seq allowed us to gain insight into how both components interact with, and respond to each other, focusing on the role of QS molecules and downstream signaling. Finally, we benchmarked our findings with a cohort of publicly available RNA-seq datasets from clinical samples of *P. aeruginosa* infected airways. Our co-culture model recapitulates metabolic aspects, CCR and nitrogen usage, as well as the expression of several secretion systems, important for *P. aeruginosa* persistence and virulence. Furthermore, the upregulation of genes involved in *P. aeruginosa* antibiotic resistance could be of particular relevance for research of bacterial mechanisms of antibiotic resistance and discovery of novel antibacterial compounds. This is highly relevant in the case of *P. aeruginosa* due to its high intrinsic resistance^11^. Since it is not possible to study early stages of infection using clinical isolates, our model offers a tool to understand the initial steps of the infectious process in near-physiological conditions.

Previous attempts to co-culture *P. aeruginosa* in 2D have been performed using cancer cell lines^32–36,78,79^ or primary human airway cultures^34,37^. The latter offers clear advantages over the former, because of the non-cancerous nature of the primary cultures. Human airway organoids allow for the indefinite biomass expansion and thus for longitudinal experiments using a defined and constant organoid source. Additionally, the indefinite expansion of airway organoids will enable *P. aeruginosa* co-culture with genetically engineered organoid lines and isogenic WT controls, once genome editing of airway organoids becomes efficient enough to perform experiments at this scale^80^. This will open the door to understanding which epithelial factors shape the course of *P. aeruginosa* infection and how to harness them to fight the infection.

In this dataset, we do not observe a major organoid response specific to QS pathways. Using live co-cultures could lead to lower effective concentrations of QS-induced molecules, compared to what has been used in studies testing the effect of single QS compounds on epithelial cells^81,82^. In addition, the strong LPS-induced inflammation, present in all conditions, might abrogate the effect of QS-derived molecules. Particularly, LPS is not accounted for in studies that solely focus on QS-derived molecules. Additionally, it is likely that only specific cell types respond to QS molecules, i.e. chemosensory tuft cells^83,84^, and therefore bulk RNAseq would not allow the study of these cell-specific effects.

Future expansion of the co-culture infection models, with the addition of immune cells, will yield insight into how this important aspect affects the behavior of the *P. aeruginosa* infections. Importantly, co-culture models that recapitulate a more complex tissue architecture^85^ will also help to understand biofilm formation under more physiological conditions. Furthermore, QS pathways coordinate the population-scale behavior of individual bacteria, which leads to the functional and spatial heterogeneity found in bacterial biofilms. The recent application of spatial transcriptomics to *P. aeruginosa* biofilms grown on solid surfaces have allowed detailed study of these two aspects^86,87^. It will be very interesting to address this spatial and functional heterogeneity in the presence of the epithelial and immune cells using co-cultures. Furthermore, our 14-hour co-culture system represents an early stage of the infectious process. While this time span allows studying QS regulation, it is too short to focus on biofilm development and other aspects of chronic infections. Longer incubation using our method was technically challenging due epithelial damage caused by the bacterial cells. Developing co-culture strategies that enable sustained chronic infection of the mucosa will help to investigate these aspects of later infection stages.

In conclusion, 2D organoid co-cultures with *P. aeruginosa* represent a new development of the current methods to study host-bacterium interplay. The system recapitulates major infection traits from both bacteria and epithelium, including bacterial metabolism, expression of virulence factors and the induction of an inflammatory response in the epithelium.

## Acknowledgements

We acknowledge the Utrecht Sequencing Facility (USEQ) for providing sequencing service and data. USEQ is subsidized by the University Medical Center Utrecht and The Netherlands X-omics Initiative (NWO project 184.034.019). We would also like to thank Dr. Tim Holm Jakobsen for kindly providing PAO1-GFP strain, Dr. Bart Bardoel for kindly providing *E. coli* RHO3 strain, and Joe J. Harrison for kindly providing the pEX18Gm plasmid.

