## Supplemental Material for "Establishment and characterization of a new *Pseudomonas aeruginosa* infection model using 2D airway organoids and dual RNA sequencing"

**Supplementary table 1 (Primers used)**

| Primer name | Primer sequence |
|---|---|
| *lasI*.UP.Fw | GCCAGTGCCAAGCTTGCATGCGAGGCCAACCGTTTCATG |
| *lasI*.UP.Rv | GCTGTTCCACCAGTACGATCATCTTCACTTCCTCCAA |
| *lasI*.DN.Fw | GAAGATGATCGTACTGGTGGAACAGCGACTGG |
| *lasI*.DN.Rv | CATGATTACGAATTCGAGCTTTCCTGCCCTGGATAGAAC |
| *lasI*.seq.Fw | GCTCGGAAGCCAATGTGAACTT |
| *lasI*.seq.Rv | AACTGGAACGCCTCAGCCAG |
| *rhlI*.UP.Fw | CATGATTACGAATTCGAGCTCGACCAGCAGAACATCTC |
| *rhlI*.UP.Rv | TGAAGCTAATTCGATCATGCATGAGCTCCAGCGATTCAGAGAGCAA |
| *rhlI*.DN.Fw | ATCCCCAATTCGATCGTCCGGCTACCACCCGGAATGGCT |
| *rhlI*.DN.Rv | GCCAGTGCCAAGCTTGCATGCCAGGTTGATCGAGATGC |
| *rhlI*.seq.Fw | ATGTCCTCCGACTGAGAGGG |
| *rhlI*.seq.Rv | CAGAGAGACTACGCAAGTCGG |

**Supplementary table 2 (Plasmids used)**

| Plasmid | Use | Reference |
|---|---|---|
| pEX18Gm | Backbone plasmid used for the generation of deletion constructs | Hmelo (2015)[45] |
| pEX18Gm::Δ*lasI* | Plasmid containing *lasI* deletion construct for use in PAO1 | This study |
| pEX18Gm::Δ*rhlI* | Plasmid containing *rhlI* deletion construct for use in PAO1 | This study |

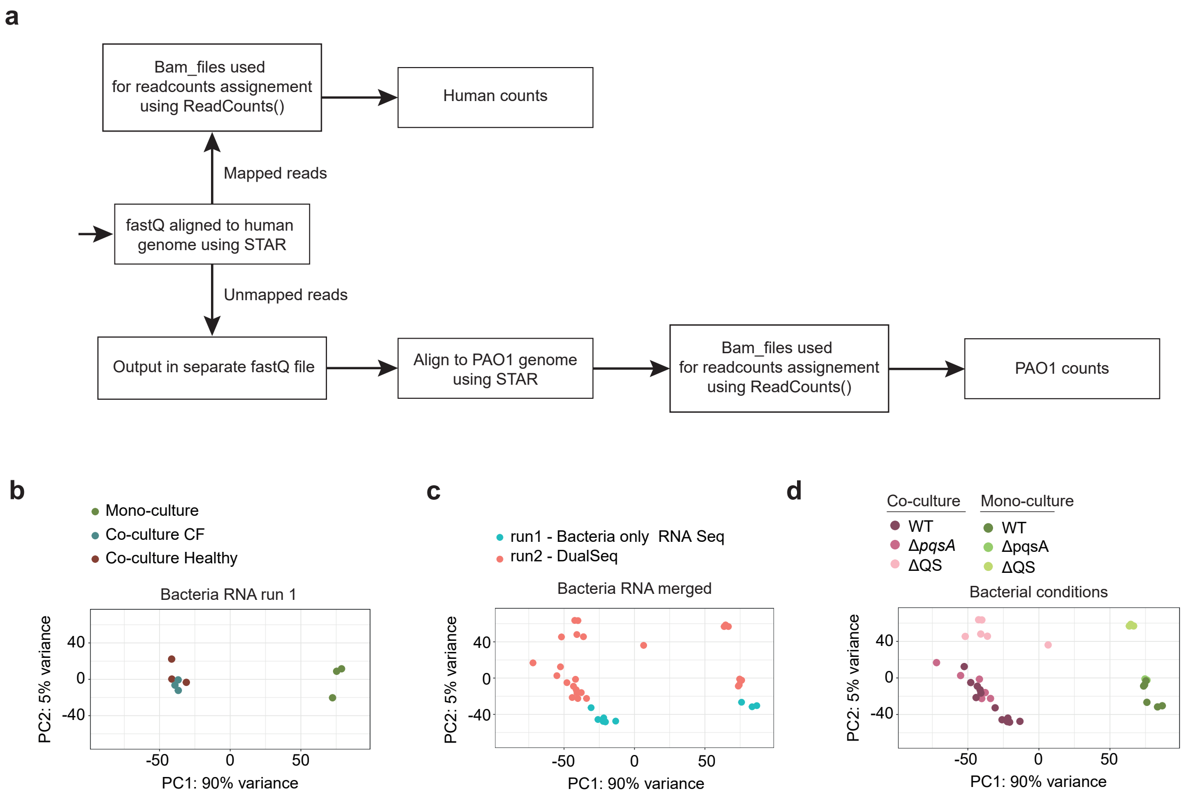


**Supplementary figure 1. Mapping strategy and PAO1 bulk dataset integration. a)** Mapping and count assignment strategy. **b)** PCA plot of PAO1-only bulk RNA samples from run 1. **c)** PCA plot showing samples by run (PAO1-only bulk RNA-seq or Dual RNA-seq). **d)** PCA plot of the integrated dataset color-coded by culture type and PAO1 genotype.

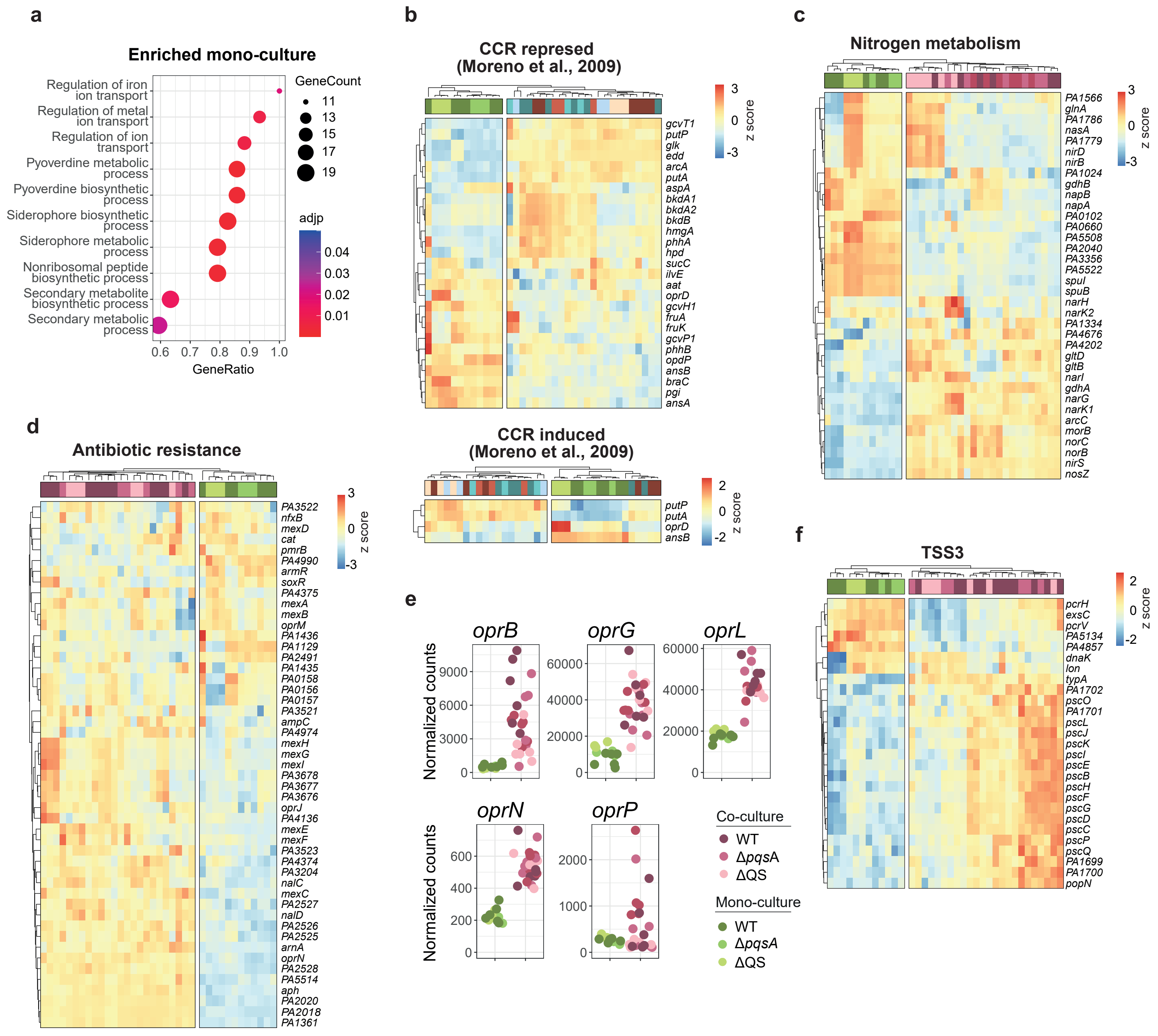


**Supplementary figure 2. Extended transcriptional response of PAO1 to the presence of airway epithelium. a)** Gene ontology enrichment analysis showing top 10 categories enriched in PAO1 mono-culture **b)** Expression heat map of genes regulated by the Carbon catabolite repression (CCR) pathway in the related bacterium *Pseudomonas putida* (Moreno et al., 2009[88]). **c)** Expression heat map of genes from the KEGG pathway nitrogen metabolism (pae00910). **d)** Expression heat map of genes known to confer antibiotic resistance to *P. aeruginosa*. **e)** Normalized count plots of DEGs encoding *P. aeruginosa* porins. **f)** Expression heat map of T3SS genes (GO:0030254).

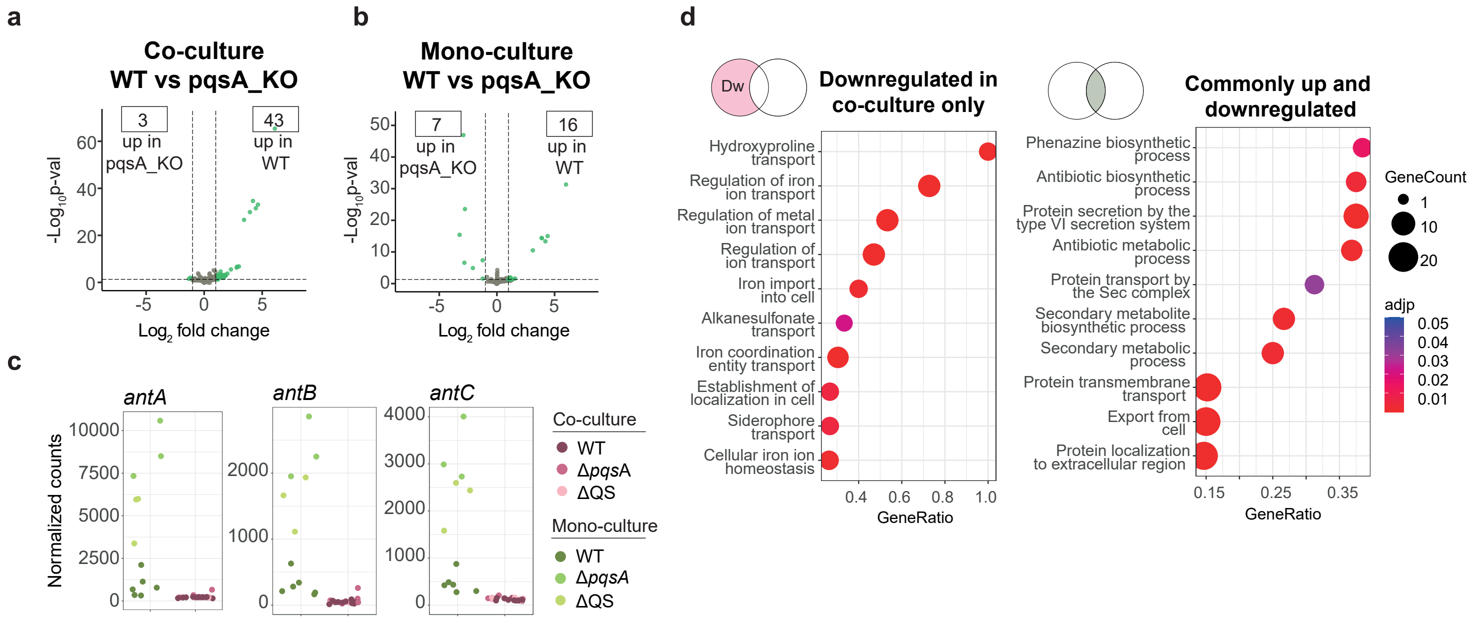


**Supplementary figure 3. Extended effects of the epithelium on PAO1 QS regulation.** **a)** Volcano plot displaying gene $\log_2$ fold change and $-\log_{10}$ adjusted p-value when comparing the transcriptomes of WT to Δ*pqsA* PAO1 in co-culture. **b)** Volcano plot displaying gene $\log_2$ fold change and $-\log_{10}$ adjusted p-value when comparing the transcriptomes of WT to Δ *pqsA* PAO1 in mono-culture. **c)** Normalized count plots of genes from the anthranillic acid metabolic pathway. **d)** Gene ontology enrichment analysis showing top 10 categories enriched in genes that are specifically downregulated in co-culture in WT and Δ*pqsA* PAO1 transcriptomes compared to ΔQS (left) or those that are common to both (right, up and downregulated).

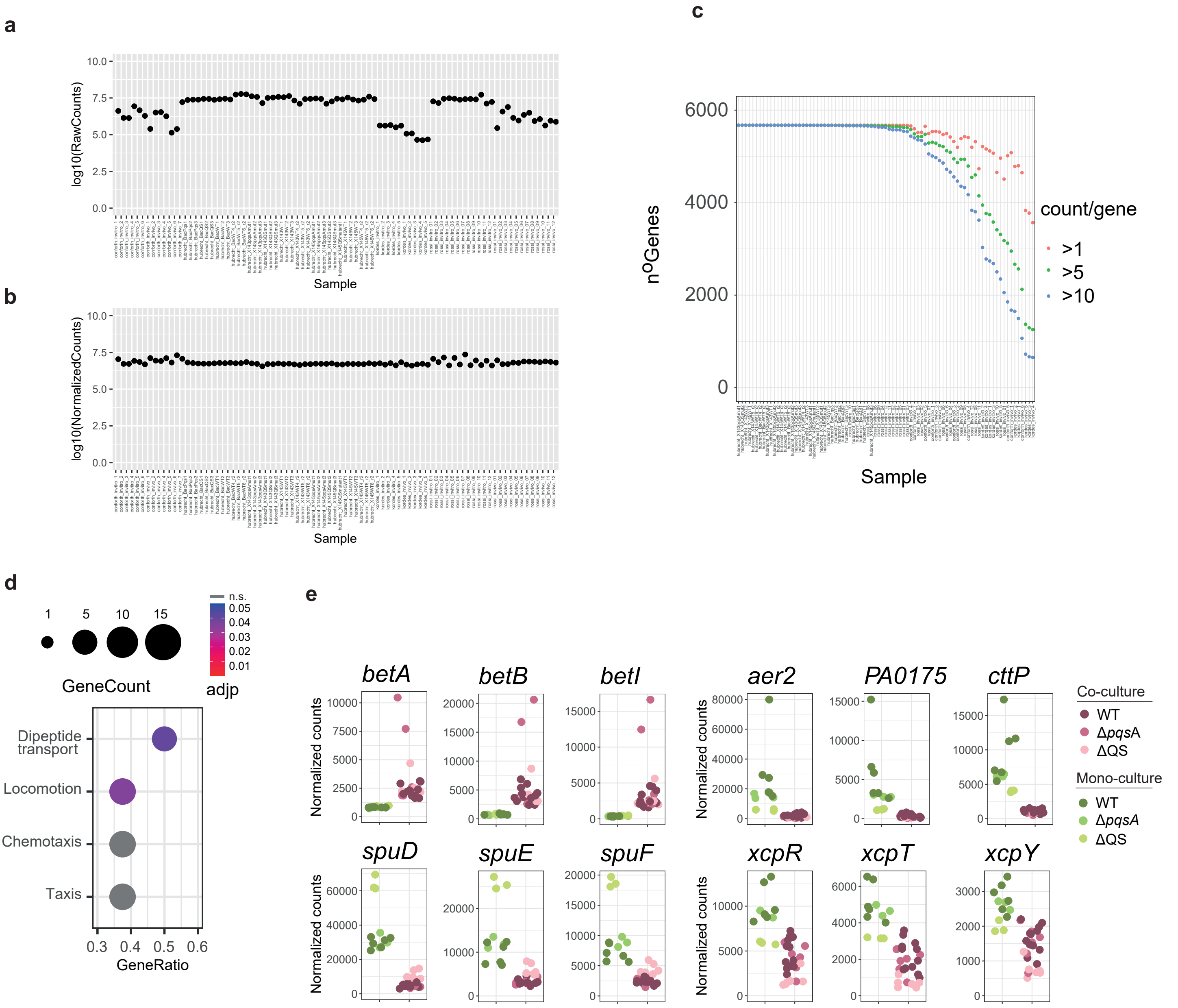


**Supplementary figure 4. Cohort integration quality control. a)** Log10 of total raw counts per sample before DESeq2 normalization. **b)** Log10 of normalized counts per sample after DESeq2 normalization. **c)** Number of genes with more than 1 (red), 5 (green) or 10 (blue) counts per sample. **d)** Gene ontology enrichment analysis showing categories enriched in top common DEGs from co-culture and *in vivo* samples. **e)** Normalized count plots of genes from Figure 6 f, performing the analysis only in the samples from our cohort, to exclude bias in the results due to the integration process with the extra datasets.

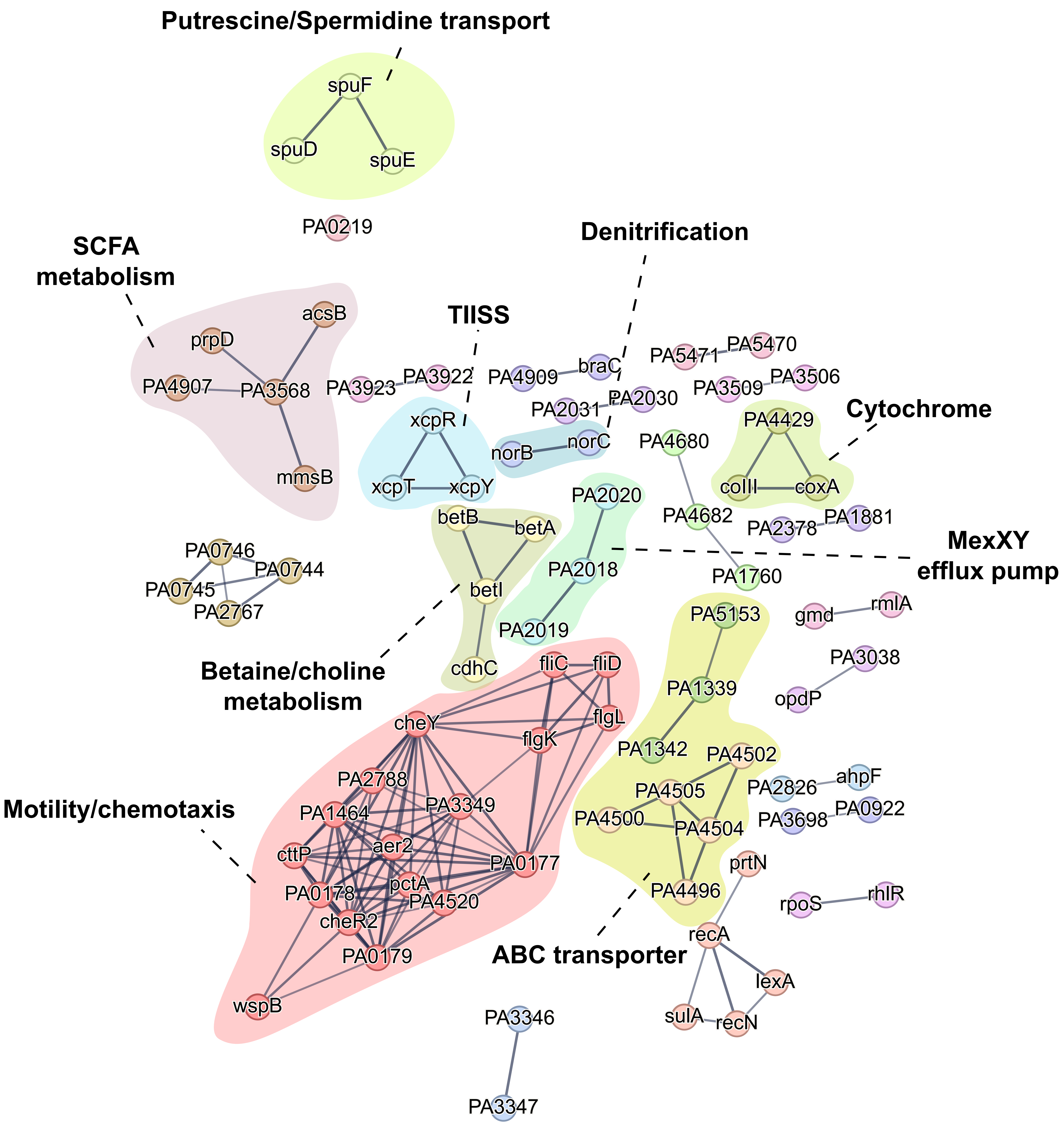


**Supplementary figure 5. Extended protein-protein network results.** Protein-protein interaction network of common DEGs (*in vivo* and co-culture). Each node represents a protein encoded by DEG. Individual gene names are highlighted over the nodes. Edges represent known protein-protein association (either physical or functional) with a confidence level higher than 0.7. Node color represent clusters generated MCL method. Highlighted the pathway to which the cluster proteins belong.
